# Aird-ComboComp: A combinable compressor framework with a dynamic-decider for lossy mass spectrometry data compression

**DOI:** 10.1101/2023.05.04.539411

**Authors:** Miaoshan Lu, Junjie Tong, Ruimin Wang, Shaowei An, Jinyin Wang, Changbin Yu

**Affiliations:** Zhejiang University, Hangzhou 310009, Zhejiang Province, China; School of Engineering, Westlake University, 18 Shilongshan Road, Hangzhou 310024, Zhejiang Province, China; Institute of Advanced Technology, Westlake Institute for Advanced Study, 18 Shilongshan Road, Hangzhou 310024, Zhejiang Province, China; Shandong First Medical University, Jinan 250000, China; Fudan University, Shanghai, China; School of Life Sciences, Westlake University, 18 Shilongshan Road, Hangzhou 310024, Zhejiang Province, China; Institute of Biology, Westlake Institute for Advanced Study, 18 Shilongshan Road, Hangzhou 310024, Zhejiang Province, China

**Author notes:** These authors contributed equally to this work (Miaoshan Lu, Junjie Tong).

## Abstract

Mass spectrum (MS) data volumes increase with an improved ion acquisition ratio and a highly accurate mass spectrometer. However, the most widely used data format, mzML, does not take advantage of compression methods and improved read performances. Several compression algorithms have been proposed in recent years, and they consider a number of factors, including, numerical precision, metadata read strategies and the compression performance. Due to limited compression ratio, the high-throughput MS data format is still quite large. High bandwidth and memory requirements severely limit the applicability of MS data analysis in cloud and mobile computing. ComboComp is a comprehensive improvement to the Aird data format. Instead of using the general-purpose compressor directly, ComboComp uses two integer-purpose compressors and four general-purpose compressors, and obtains the best compression combination with a dynamic decider, achieving the most balanced compression ratio among all the numerous varieties of compressors. ComboComp supports a seamless extension of the new integer and generic compressors, making it an evolving compression framework. The improvement of compression rate and decoding speed greatly reduces the cost of data exchange and real-time decompression, and effectively reduces the hardware requirements of MS data analysis. Analyzing mass spectrum data on IoT devices can be useful in real-time quality control, decentralized analysis, collaborative auditing, and other scenarios. We tested ComboComp on 11 datasets generated by commonly used MS instruments. Compared with Aird-ZDPD, the compression size can be reduced by an average of 12.9%. The decompression speed is increased by an average of 27.1%. The average compression time is almost the same as that of ZDPD. The high compression rate and decoding speed make the Aird format effective for data analysis on small memory devices. This will enable MS data to be processed normally even on IoT devices in the future. We provide SDKs in three languages, Java, C# and Python, which offer optimized interfaces for the various acquisition modes. All the SDKs can be found on Github: https://github.com/CSi-Studio/Aird-SDK.

## Introduction

With the improvement of the accuracy of mass spectrometers and the optimization of acquisition methods, the ion signals collected by mass spectrometers have become much more abundant, resulting in an explosive increase in MS data files. Finding better ways to compress MS data files while maintaining high read and write performances is a much-needed advancement. As one of the pervasive data formats, mzML is still the preferred format for many laboratories.^1^ By using Proteomics Standards Initiative-Mass Spectrometry (PSI-MS) controlled vocabulary (CV) with Extensible Markup Language (XML),^2^ mzML can completely record almost all the metadata in the original vendor file with a human-readable format. Due to the mixed storage of the base64-transcoded spectral data and metadata, the compression ratio is not impressive, which makes mzML appear bloated. Using tools such as MSConvert^3^ can convert vendor files to mzML or other open formats but there is no way to convert mzML back to the vendor files. Vendor files also have a high compression ratio, so most laboratories use vendor files as the main archive format. However, the vendor format also has limitations. Reading different vendor formats requires a specific additional Software Development Kit (SDK), and some of the kits do not support cross-platform file reading. In the proteomics or metabolomics analysis workflow, the huge memory consumption and bandwidth consumption caused by large volumes complicate the omics calculations based on mass spectrometry. Therefore, it is necessary to construct a unified, computing-oriented, cross-platform reading data format.

MS data are mainly divided into binary data and metadata. Binary data consist of *m/z*, intensity, and ion mobility from Trapped Ion Mobility Spectrometry (TIMS) data,^4^ which is the largest component of the MS file. To store metadata and binary data more elegantly, separating the two parts, using cross-platform data storage files for metadata and a specific compression algorithm for binary data, have become an effective solution. Many new compression algorithms have been developed recently.^5–11^ From the perspective of the wide application and ease of use of the format, a widely used and computationally oriented data format must satisfy:

1. cross-platform metadata reading capabilities;
2. flexible and scalable indexing capabilities;
3. high-performance random accessing capabilities;
4. stable and balanced MS data compression capabilities;
5. complete SDK data reading with rich documentation;
6. basic toolset or third-party extensions.

In terms of the storage medium of the format file, mzDB^9^ uses SQLite^12^ as the file storage technology, mzDB has implemented many calculation optimizations and has created related designs for data-dependent acquisition (DDA) and data-independent acquisition (DIA) MS acquisition methods, which improves the running speed of the extracted ion chromatogram (XIC) process. However, mzDB only saves 25% of the storage space compared with mzML. mzMLb^11^ uses Hierarchical Data Format version 5 (HDF5),^13^ a widely recognized cross-platform file storage technology. HDF5 also improves the data read/write performance and compression ratio based on a rich plug-in system and cache system. Aird stores the metadata with JavaScript Object Notation (JSON), and the spectra data are additionally stored as binary data so researchers can quickly preview the metadata of the mass spectrum file. Work in Aird-ZDPD shows that using JSON instead of XML to store the same metadata information can reduce the volume by an average of 53%.^5^ Since JSON is widely compatible with browsers and some text editors, it is read-friendly in most scenarios.

In terms of the numerical precision and compression ratio, StackZDPD, ^6^ mspack^8^ and Numpress^14^ stipulates the *m/z* precision loss in detail. mspack specifies the numerical precision that the computer can retain under a 32/64-bit floating point. StackZDPD shows that, compared with mzML, using six decimal places (6dp, 0.002ppm at 500 daltons, similar conclusions are also verified in the manuscript of Numpress and mzMLb) for *m/z* would not affect the downstream of a given workflow of an ADAP chromatogram building on MZmine.^15^ StackZDPD also tested the impact of downstream calculations in 5dp mode. Approximately 28 differential peaks were identified out of 20642 peaks.^6^ Most of the peaks belonged to some interfering peaks generated in the case of extremely small precision values. The final chromatographic identification only has one variation in the result. In a recent study, Tong et al. conducted a comprehensive experimental comparison to determine how the relative accuracy of the numerical values affected the outcomes of the mass spectral data recognition. The outcome indicates that the intensity value has little bearing on the feature extraction, feature quantification, and compound identification when the relative error is 2 *×* 10*^−^*^3^. The *m/z* has negligible impact on feature extraction, feature quantification, and compound identification when the relative error is 10*^−^*^5^.^16^

In terms of the compression ratio, StackZDPD has a better compression ratio than mzMLb (mzLinear+trunc+zlib) and the ZDPD compressor. However, due to its complex encoding process, its compression and decompression speeds are lower than those of ZDPD. mspack proposes a compression method based on the similarity of the adjacent spectra, and directly uses a general-purpose compressor for the compressions after a data transformation. In contrast, StackZDPD uses the stacking spectra technique to further increase the encoding similarity of the *m/z* values. Since *m/z* is an ordered array, an encoding conversion step is first performed using methods such as the delta transform or linear prediction transform. The converted array will have a higher similarity and smaller values. Then, a general-purpose compressor based on LZ77^17^ is used for compression. Compressing too many spectra can make the random files less readable. Random file accessing is also a point to consider when designing the MS data format. More consideration must be given to the random accessing file performance if it used for the calculation process. For storage, the compression ratio is the primary consideration.

Aird is a computation-oriented format. The core compression algorithm in Aird improves the compression ratio of a single spectrum while keeping the random access capability unchanged. The original ZDPD compressor first proposed to use integers as the actual *m/z* storage value types and further reduce the number of encoded bits by using the integer compressor binary packing (BP) instead of using a direct compression with a general compressor. The basic idea of the BP is that for an integer data block, most of the data can be stored in a small space, and the remaining part is treated as abnormal data. Generally, by setting a value of the frame bit, it can store more than 90% of the data, and that part is called the normal part, and the remaining less than 10% of the data are stored separately, and is called the abnormal part. Since the BP does not change the expression mode of the same value, it does not change the compressed data similarity. Therefore, using the BP algorithm does not alter the similarity of the compressed data too much. This enables subsequent general-purpose compression algorithms to attain extremely high compression capabilities. At the same time, the decompression performance of the BP algorithm using single instruction multiple data (SIMD) acceleration technology is extremely fast.

Using an additional integer compressor proves to be an advantage in compressing a single spectrum.^5^ However, the following issues are not discussed in the work of Aird-ZDPD: (a) Expansion with more new compressors. Decompression time is as important as the compression ratio for a computationally oriented format. Is there better performance with other integer-purpose compressors and a combination of general-purpose compressors? How can a new compressor be seamlessly extended in the future without changing the schema of the original format? (b) Expand more data arrays with new dimensions. The Aird format optimizes *m/z* arrays and only uses the traditional Zlib compressor for intensity arrays. Is it possible to introduce ZDPD into the intensity dimension? What compression strategies for new dimensions may be presented in the future, such as ion mobility data from PASEF(parallel accumulation-series fragmentation). (c) Are there multilanguage SDKs, detailed description documents, a rich and complete API interface to help developers use it easily?

With these three issues in mind,we developed ComboComp with Aird, a combinable compressor framework with dynamic decisions for near-lossless MS data compression. ComboComp systematically evaluates the performance comparisons of the various compressors under the combination framework. In actual tests, we found that a single compressor cannot achieve optimal solutions on all the formats due to the different distribution characteristics of the values in the different MS files. So that the developers do not have to bear a large memory burden, ComboComp introduces a dynamic decider that can dynamically predict which combination can attain the best performance. To further improve the usability of the Aird format, we have simultaneously provided SDKs for Java, Python and C#. In addition, more interfaces are provided for the DIA, ^18^ DDA, diaPASEF^19^ and ddaPASEF^20^ acquisition modes.

ComboComp integrates the algorithm implementations of two integer-purpose compressors. In addition, it incorporates four general-purpose compressors. The float-to-integer encoder for *m/z*, intensity and ion mobility has also been implemented individually (see Figure 1). ComboComp also uses an empty integer compressor, so it can be compatible with a single general-purpose compressor. We found that in some small-size MS file compressions, using the general-purpose compressor directly can achieve a better compression performance. In any case, deciding which combination is ultimately used can be handled by the dynamic decider for a prediction.

In this work, we tested files from four manufacturers: Agilent, Bruker, Sciex and Thermo. The acquisition mode involves DDA, DIA, ddaPASEF and diaPASEF (PASEF uses aggregation mode). Compared with ZDPD, the compression ratio of ComboComp is increased by an average of 12.9%, while the decompression speed does not drop but increases by an average of 27.1%. To make the Aird format easier to use, we also extended the Aird format for the latest version of mzmine3.^21^ We tested it on an Intel i9-10gen PC with 16 GB memory and read a 600 MB vendor format file (loading all the spectra data into memory). The vendor format takes 80 seconds, the mzML takes 90 seconds and the Aird format takes only 5 seconds.

Extending MS data to cloud computing has been a difficulty in this field. The first and most important problem is that the limited bandwidth can not effectively support the large raw data. With the increasing attention to energy efficiency and performance of data transmission in the field of network computing. ^22–24^ Aird format provides a reliable and feasible way for MS files transmission on the network, which provides the foundation for cloud computing.

## Materials and Methods

### Extensible Combination Compression Framework

ComboComp builds a standard compression pipeline framework and embeds two integer-purpose compressors (binary packing and variable byte) and four general-purpose compressors (Zlib, Brotli, Snappy and Zstd) that are tested and effective.^25–28^ The framework also supports compression under a single integer or general compressor. The two integer compressor libraries come from the work of Lemire.^25^ The BP and VB algorithms are two effective integer-purpose compressors.

The actual storage length of the small integers is shortened by the variable step size and fixed step size. At the same time, the related encoding library provides the streaming SIMD extension (SSE) acceleration capability, which makes the compression and decompression speed much higher than those of similar compressors. The new method changes the limitation of the traditional data format to compress MS data by using a unique compression algorithm. After performing comprehensive tests, we found that no global optimal compression algorithm exists. Although the objects stored in the MS files are similar, the different acquisition modes and vendor files have different digital distribution characteristics. Different MS files can achieve improved compression performances by using their optimal compression schemes. Moreover, in the implementation of AirdPro, ^5^ we specifically added the full enumeration verification mode. It can generate all the combined compression results at one time, which makes it convenient for the newly expanded compression kernel to perform a comparison.

For the *m/z* arrays, eight combinations (two integers times four general compressors) are tested. However, for the remaining data, i.e., intensity and ion mobility, the empty integer compressor and the other two compressors (i.e., BP and VB) coupled with four types of general compressors (i.e., Zlib, Snappy, Brotli and Zstd), means a total of 12 combinations are used. Therefore, for the MS data in non-PASEF mode without the ion mobility data, we need 8 *×* 12 = 96 enumerations to compare all the results for each file. For the PASEF mode with the ion mobility data, we need 8 *×* 12 *×* 12 = 1152 enumerations. Since the size of the MS files is inherently large, it may take months to test and complete all the conversions. Therefore, we use the sampling method to compare each file in parallel. In this sampling strategy, we randomly pick 500 spectra from each file and triplicate this random process to obtain the average value. The data distribution characteristics of mass spectra are similar in the same file, so it is a relatively reliable process to collect 500 spectra for prediction use. We also completely transform part of the dataset and compare it with the predicted results to demonstrate this.

### Dynamic decider to predict the best combination

Since the combination framework will involve various combination methods, it is necessary to obtain the optimal solution. ComboComp presents a dynamic decider that makes predictions through simple random sampling. Part of the spectral data (default 500) is randomly selected for a compression test with all the combinations, and finally, the compressed size (referred to as *S*), compression time (referred to as *C*) and decompression time (referred to as *D*) are counted. *n* is the number of all the combination types available in the system.

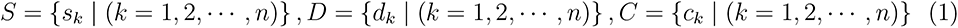

Since the units and distributions of these three data types are different, we need to perform a max-min normalization transformation before scoring.

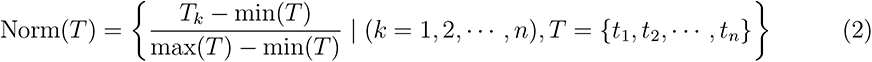

After a normalization transformation on *S*,*C*,*D*, we obtain three new arrays of *S^′^*,*C^′^*,*D^′^*.

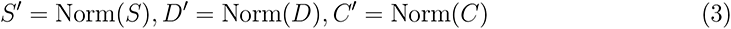

To avoid the compression ratio being too low, we calculate the mean(*S^′^*) of *S^′^*. A threshold of mean(*S^′^*) would be delineated as a baseline. The compressed file size must be under the baseline. For a compliant combo compressor in *S^′^*,*C^′^*,*D^′^*, we add them with a weight of a:b:c (default is 1:1:1) to obtain a new array. Users can modify this weight ratio according to their needs. The smallest value in this array is our final predicted combination:

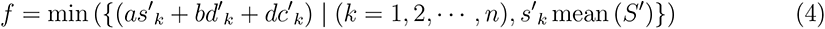

Using the decider for dynamic selection of the optimal combination can effectively improve the performance of the compressors on different datasets. Although the compression principle of each compressor will not change much in the future, their implementation algorithms may be continuously optimized. ComboComp will inevitably introduce better compressors or upgrade existing compressors to as part of its evolution.

### Compression method for intensity arrays

An intensity array is a set of data with widely varying characteristics. Intensity values exhibit huge repeatability on some instruments, especially at low concentrations. Thus, the result will show many repeating low signals, enabling a general-purpose compressor to have a high compression ratio. However, for some instruments, its values present a periodic normal distribution, with weights ranging from 0 to hundreds of millions, which greatly reduces the effectiveness of the general-purpose compressors. Introducing ComboComp into the intensity array compression process is worth trying. However, unlike *m/z*, the intensity array has a wide range of values, and converting the double-type intensity value to an integer type may lead to a numerical overflow. Many formats use *log* transcoding, or are accurate to 10*^−^*^4^ for lossy compression. For ComboComp, we only use *log* transcoding for numbers larger than 2^30^ *−* 1. For smaller numbers, all the significant digits are kept intact. After the log conversion, we use the sign bit of the integer to mark it. The conversion formula is as follows:

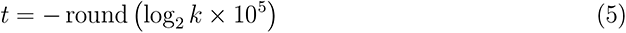

We can convert these extra large values into a 6 or 7 bit integer and then negate it as a converted value so that it can be quickly identified when decoding.

Since the maximum difference generated by rounding is 0.5 by multiplying by 10^5^, the maximum error we can control is:

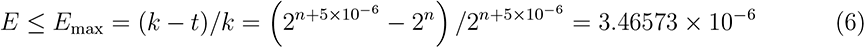

The maximum absolute error is less than 3.46573 *×* 10*^−^*^6^, which is a reasonable lossy data transformation for intensity.

### Compression method for mobility

ComboComp uses a lossless compression strategy for ion mobility. Ion mobility is an additional dimension of the data generated by the instruments that use ion mobility. It should be noted that ComboComp currently only supports data in the combined mode of Bruker TimsTOF. The ion mobility array is a set of double-type collections. On the Bruker TimsTOF instrument, the mobility array can be calculated by calling the SDK (timsdata.dll, file version: 2.7.103.49-68-vc141) provided by Bruker with a given index range. Therefore, the mobility array of the double type can be reversely mapped to various index integers. The final stored ion mobility value is lossless.

### Empty Integer Compressor

An empty integer compressor is used before the general compressor to ensure the consistency of the system conversion process. This empty integer compressor is essentially indicating that only the general-purpose compressor is used. For example, Empty-Zlib represents the generation of compression results using the Zlib compressor merely. This mode is used to demonstrate the effectiveness of using a combined compressor for intensity and ion mobility.

### Use Aird as the final converted file format

ComboComp uses Aird as the final format. Aird^5^ is a computationally-oriented, precision-controllable mass spectrometry data format. Aird uses JSON instead of XML to store the necessary controlled vocabulary metadata information. Beyond that, Aird makes logically related spectra closer by aggregating and reordering spectra for different acquisition modes in terms of the spectra storage structure. For example, two temporally adjacent spectra in DIA mode are not logically related, and it is necessary to find the same precursor spectra to establish a logical relationship. It will become more convenient and fast for data reading after reordering. Aird uses the differential encoding and combined compression method of Binary Packing and Zlib(i.e., ZDPD) for *m/z* arrays, reaching improvements in compression performance.

## Results and discussion

We performed extensive tests on 11 data files generated by instruments from four manufacturers, including Sciex, Thermo, Agilent and Bruker, and by four acquisition modes, DDA, DIA, ddaPASEF and diaPASEF(see Table 1). The size of these vendor files ranges from 120 megabytes to 21 gigabytes. Aird-ZDPD discusses the effectiveness of the combined compression workflow for *m/z* ratio compression in detail, To have a clearer understanding of whether the combined compression framework is effective in intensity and ion mobility dimensions, we compared the compression effects separately.

**Table 1:**
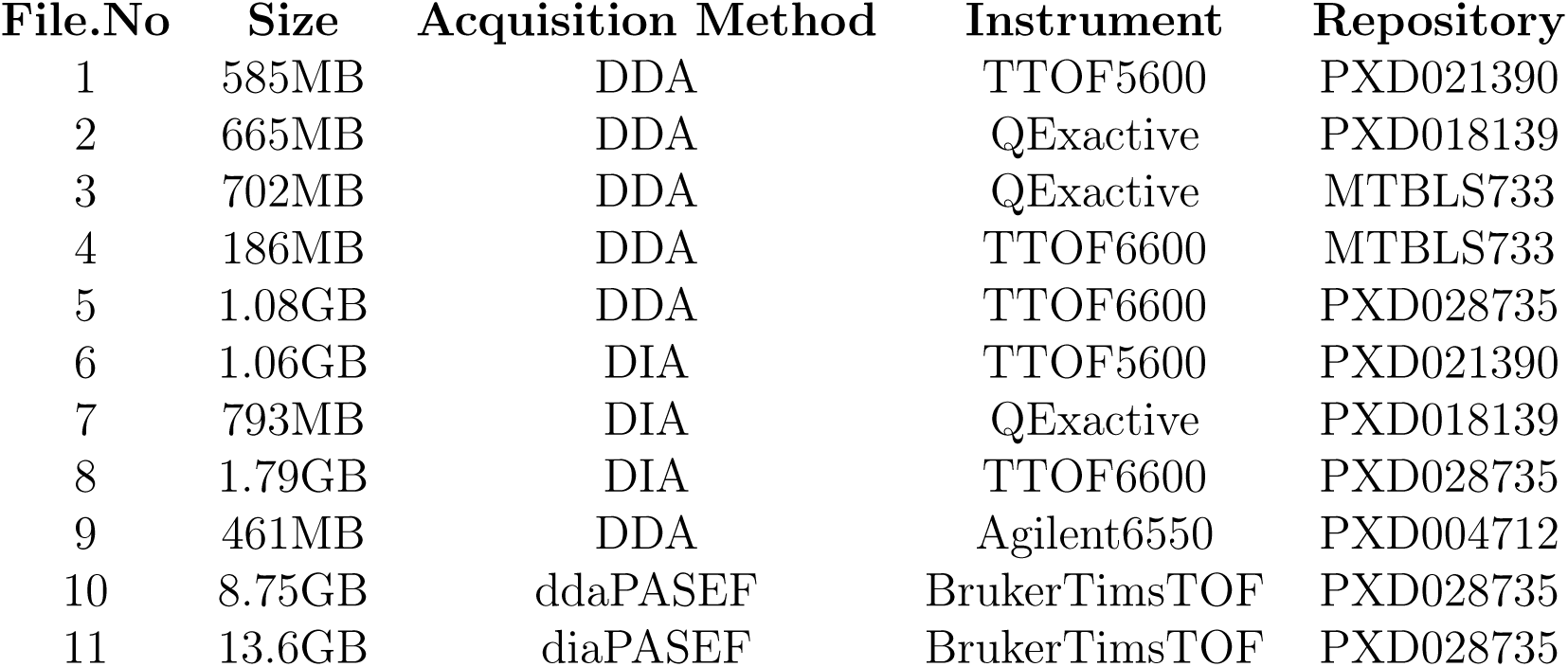
11 test files.

| File.No | Size | Acquisition Method | Instrument | Repository |
| --- | --- | --- | --- | --- |
| 1 | 585MB | DDA | TTOF5600 | PXD021390 |
| 2 | 665MB | DDA | QExactive | PXD018139 |
| 3 | 702MB | DDA | QExactive | MTBLS733 |
| 4 | 186MB | DDA | TTOF6600 | MTBLS733 |
| 5 | 1.08GB | DDA | TTOF6600 | PXD028735 |
| 6 | 1.06GB | DIA | TTOF5600 | PXD021390 |
| 7 | 793MB | DIA | QExactive | PXD018139 |
| 8 | 1.79GB | DIA | TTOF6600 | PXD028735 |
| 9 | 461MB | DDA | Agilent6550 | PXD004712 |
| 10 | 8.75GB | ddaPASEF | BrukerTimsTOF | PXD028735 |
| 11 | 13.6GB | diaPASEF | BrukerTimsTOF | PXD028735 |

### Compression performance for intensity and ion mobility

Since combined compression is applied to the intensity and ion mobility dimensions for the first time, we conduct a detailed comparison of the compression size, compression time (CT) and decompression time (DT) between ComboComp, and only use each general-purpose compressor. The effect of using combined compressors in the *m/z* dimension has been discussed in Aird-ZDPD and achieved high compression performance.^5^ For the intensity dimension,see Figure 2-A, the results show that ComboComp also has a better compression ratio than only using the general-purpose compressor (Brotli, Zlib, Zstd and Snappy), decreasing the intensity size by 12.1%, 19.2%, 19.5% and 42.8%, respectively, As a computationally oriented format, the decompression time is an important consideration. ComboComp has obvious advantages in decompression time, the decompression time is reduced by 25.9%, 43.7%, 18.2% and 31.2%, respectively.

**Figure 1:**
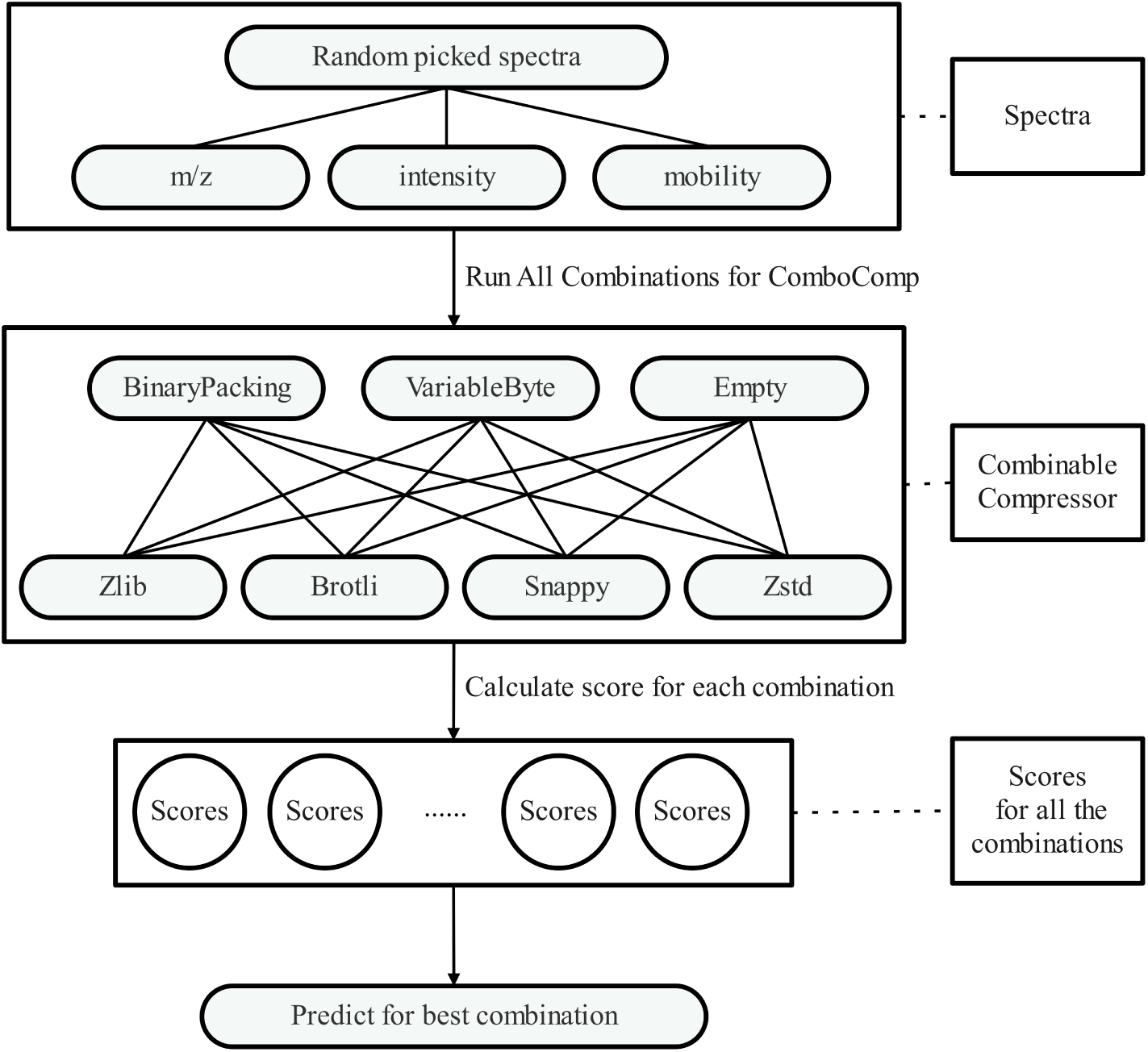
for ComboComp. We randomly pick multiple spectra and test the arrays composed of three-dimensional data, i.e., *m/z*, intensity and mobility (if exists). We then convert each data point into an integer array and compress it via eight combinations of combined compressors (two integer compressors times four general compressors). Each combination’s result are scored from three aspects, including the compression ratio, compression time and decompression time. Finally, all the scores are processed by the function f, and the combination with the highest score is selected as this file’s final scheme.

**Figure 2:**
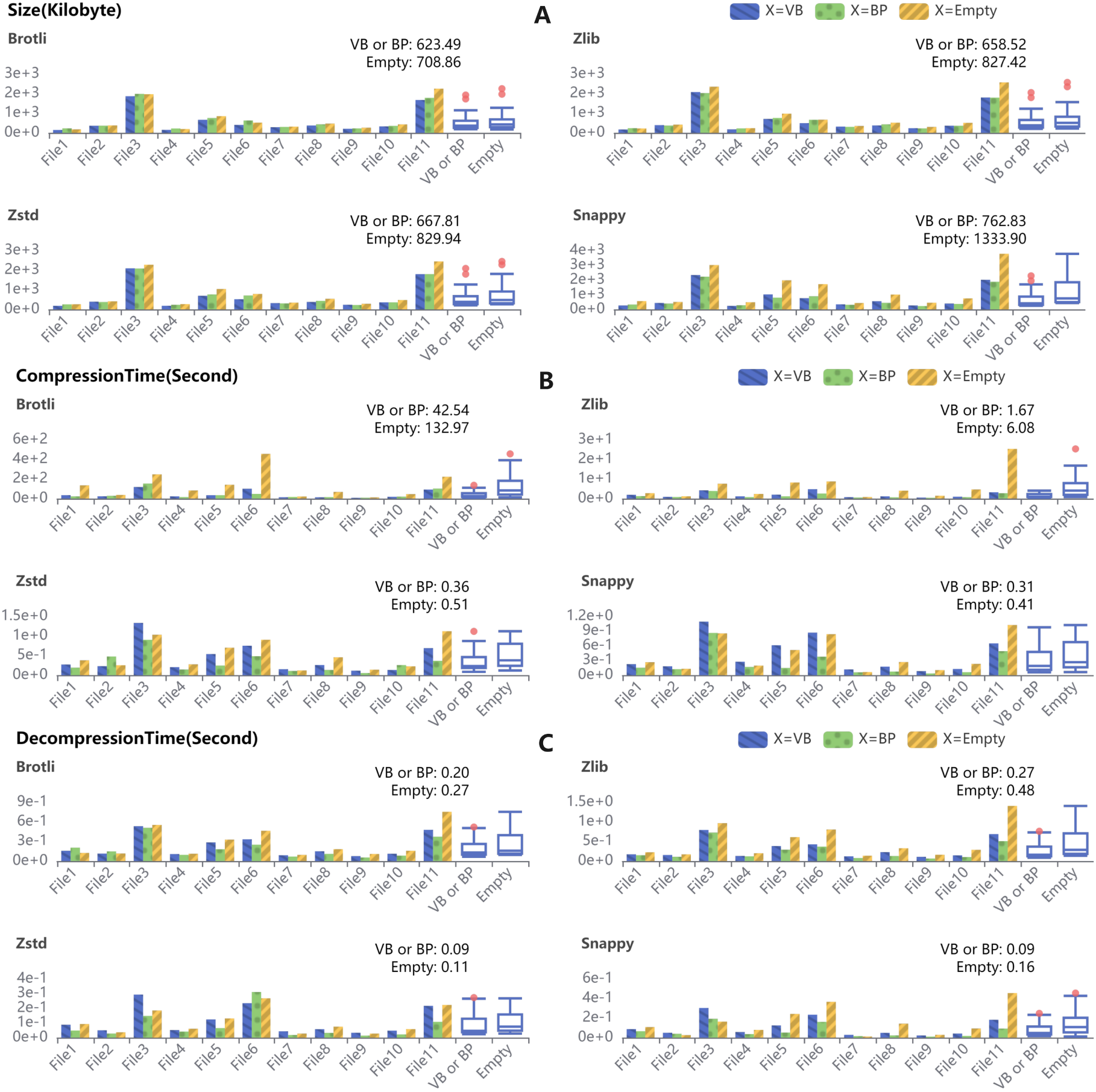
Comparision of compression size, compression time and decompression time of ComboComp and only general-purpose compressor on the intensity. We randomly collect 100 spectra from 11 test files over three times. (A). ComboComp decreased compression size by 12.1%, 19.2%, 19.5%, and 42.8% on Brotli, Zlib, Zstd and Snappy, respectively. (B). ComboComp decreased compression time by 68%, 72.5%, 29.4%, and 24.4% on Brotli, Zlib, Zstd and Snappy, respectively. (C). ComboComp decreased decompression time by 25.9%, 43.7%, 18.2%, and 31.2% on Brotli, Zlib, Zstd and Snappy, respectively. Comparing each type of ComboComp, we found that Zstd and Snappy involved combinations cost the least decompression time.

For ion mobility,see Figure 3-A, ComboComp decreased the ion mobility size by 6.3%, 18%, 15.7% and 27.9%, respectively. ComboComp achieved a huge improvement. and for decompression time, it is reduced by 45.2%, 77.7%, 15.1%, and 24.1%, respectively.

**Figure 3:**
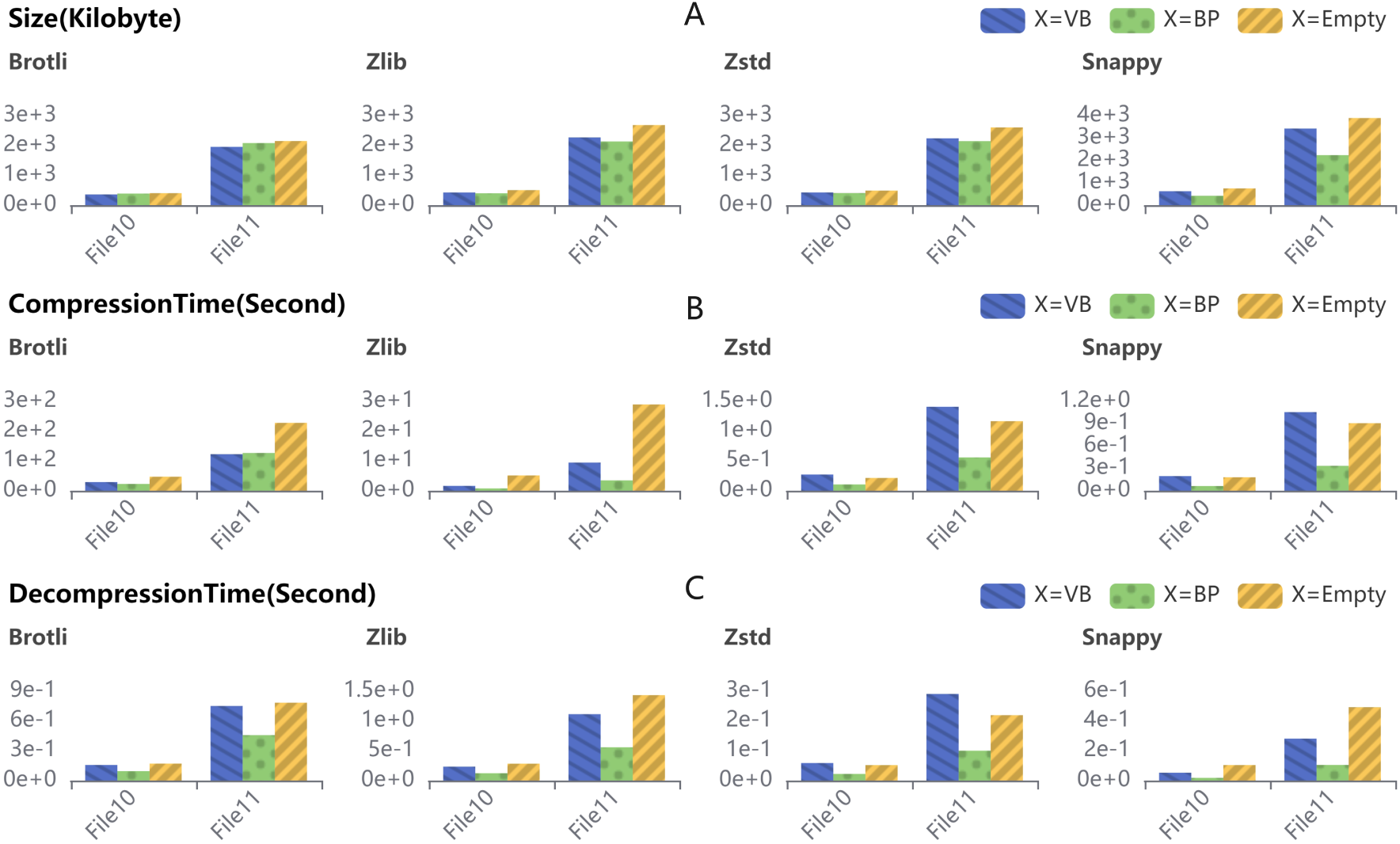
Comparision of compression size, compression time and decompression time of ComboComp and only general-purpose compressor on the mobility. Randomly collect 100 spectra over three times from two PASEF-related files. (A).Compression result on mobility is not as obvious as intensity, but still achieve a compression advantage of 6.3%, 18%, 15.7% and 27.9% in Brotli, Zlib, Zstd and Snappy, respectively. (B).ComboComp has certain advantages in terms of compression time, with a decrease of 23.6%, 40.9%, 13%, 61.8% on Brotli, Zlib, Zstd and Snappy, respectively. (C).ComboComp has great advantages in decompression time over all general-purpose compressors. the decompression time is reduced by 45.2%, 77.7%, 15.1%, and 24.1% on Brotli, Zlib, Zstd and Snappy, respectively.

### Dynamic decider process and effects

We tested the dynamic decider on 11 MS-based files(see Table 1) to see if the predicted combination is the best. Since the different combinations have different effects on *m/z*, intensity and ion mobility, we make predictions for the three data dimensions separately to obtain the best combination scheme. In Aird-ZDPD, the combination scheme of *m/z* is a fixed IBP+Zlib, and the intensity is directly performed with Zlib. We highlight these old schemes in the figure with the marker line “ZDPD”. Furthermore, we use the marker line “Selected” to mark the finalized selected combination by the decider(see Figure 6, Figure 7). The compression performance is jointly determined by three parameters, compression size, compression time and decompression time. To compare the performance of each combined compressor, we normalized the three parameters to the value *T otal*(*T*) with the dynamic-decision method. Then, we termed the *total*(*T*) as the Y-axis (smaller is better) and the combined compressors as the X-axis. We set two criteria in selecting the optimally combined compressors: (a) The compressed file *size*(*S*) must be smaller than the mean compressed file size; (b) the *total*(*T*) value is minimized while the first criterion is met.

As a result, (a) the selected optimal combination performs better than the ZDPD in all files; (b) the combined compressors IVB-Zstd and BP-Zstd are most frequently used in *m/z* and intensity, respectively, however, the other general-purpose compressors also have the opportunity to be a part of the optimally combined compressors.

Overall, for *m/z* compression, the combination of IVB-Zstd was selected as the optimal one by over 90% of the files. For intensity compression, BP was chosen as the optimal integer-purpose compressor for most of the files (9 out of 11 files, except files 4 and 6), of which seven files (files 1, 2, 3, 5, 7, 10 and 11) selected Zstd as the optimal followed by the byte compressor, and the remaining two files (files 8 and 9) selected Snappy to couple with BP. The remaining files (files 4 and 6) chose VB-Zstd as the optimal combination compressor. In each file, we compared the “Selected” with “ZDPD” for *m/z* and intensity and explained how the “Selected” meets the requirements for the optimally combined compressors.

### Converted File Comparison

We transformed 11 test files using the predicted compression combinations, measured their compression size, compression time and decompression time and compared the results with ZDPD and the vendor files. In general, compared with ZDPD, ComboComp can decrease the compression file size by an average of 12.9% (see Figure 4-A). The compression time is reduced by an average of 4.2% (see Figure 4-B). More importantly, ComboComp can relatively shorten the decompression time across all the files with an average of 27.1% (see Figure 4-C).

**Figure 4:**
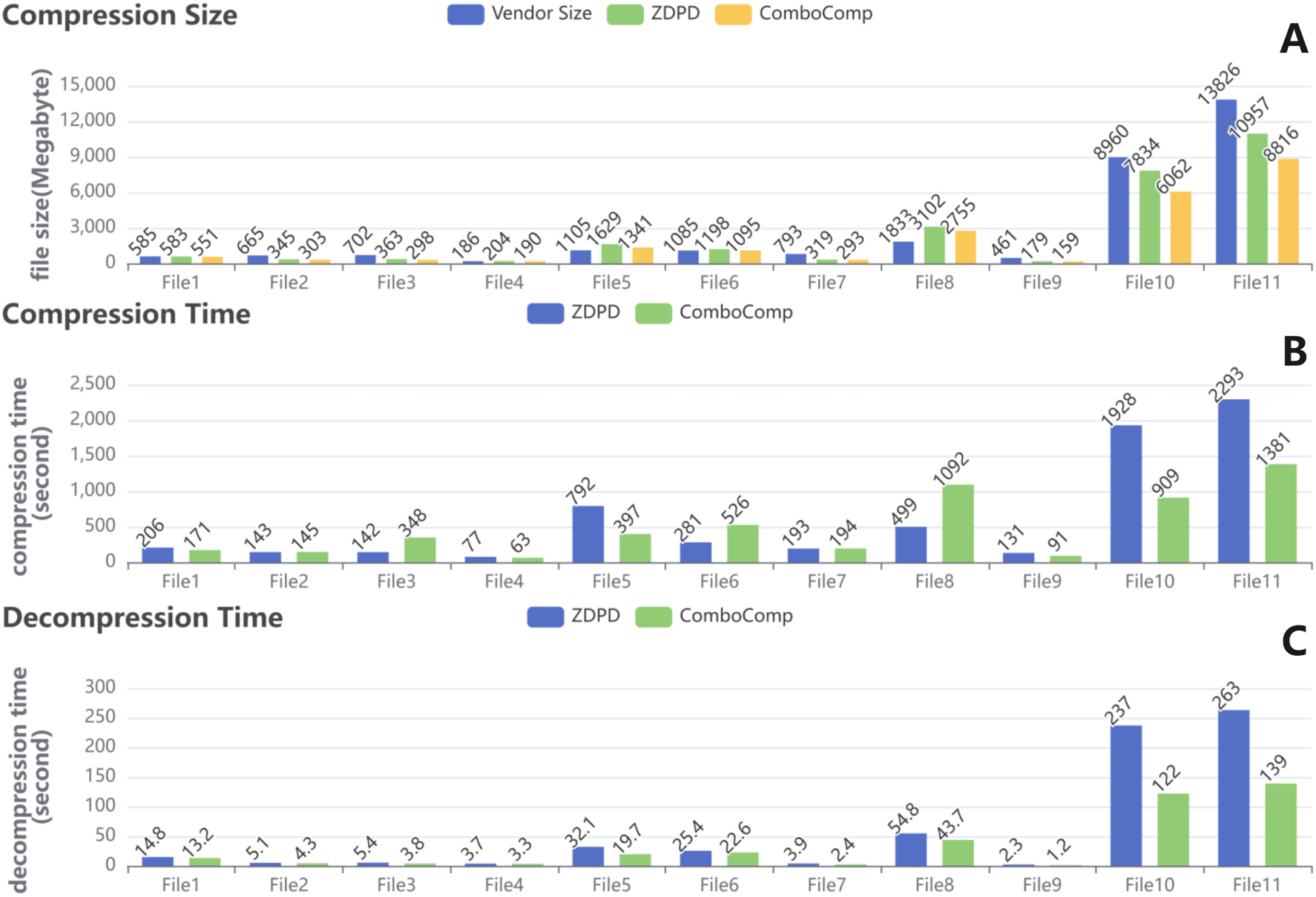
(A). Compression size comparisons of the converted files. Here, we collected and compared all file sizes across the vendor files, ZDPD and the ComboComp compressed files. (B) Compression time comparisons of the converted files between ZDPD and ComboComp. The 11 files were compressed by either ZDPD or ComboComp for comparisons of the compression times. ComboComp exhibits less compression time among these files, by approximately two times in comparison to ZDPD in all the PASEF-related data (File10: 52.9%; File11: 39.8%) and DDA-related File5 (49.9%). However, on the opposite side, ComboComp may also expend more compression time on File3 (DDA-related), File6 (DIA-related) and File8 (DIA-related). For the remaining files, a similar performance exists between ZDPD and ComboComp. (C). Comparison of decompression times between the ZDPD and ComboComp converted files. In addition to the compression times and sizes, the decompression times were also tested for all 11 files processed by ZDPD and ComboComp. Compared with ZDPD, ComboComp can relatively shorten the decompression time across all the files by up to over 45% (File9: 47.8%; File10: 48.5%; File11: 47.1%). Four of the rest of the files showed a decompression time decrement in ComboComp by over 20%, (29.6% of File3, 22.4% of File6, 38.4% of File7 and 20.3% of File8). The remaining four files compressed by ComboComp can improve the decompression speed over ZDPD files by over 10% (File1: 10.8%; File2: 15.6%; File5: 10.8%), except for 6.9% in File4.

In summary, ComboComp retains a generally faster decompression speed than ZDPD among all the files over time, with an average of 27.1%. Furthermore, the file size positively correlates with the decompression speed improvements. As we discussed above, the compression size and decompression time tend to matter more than the compression time because one file is usually compressed once but is decompressed multiple times.

As a new lossy compressor, mspack has a high compression ratio. mspack is designed for storage and currently has no random accessing capabilities. For a more comprehensive compressed volume comparison, we converted the test data sets to mz5 with 32-bit for *m/z* and intensity, to mzML with 32-bit for *m/z* and intensity, to mzMLb with “–mzTruncation=19 –intenTruncation=7”, and to mspack with default loss parameter. Since some of the format cannot compress the format in PASEF mode, we only used File1-File9 for testing. Same as the conclusion of Aird-StackZDPD and Aird-ZDPD, Aird-ComboComp has great compression advantages compared with mz5, mzML and mzMLb. For mspack, the results show that ComboComp has a better compression ratio in five of the nine tested files, but mspack achieves a clearer compression advantage in the remaining three files (see Figure 5).

**Figure 5:**
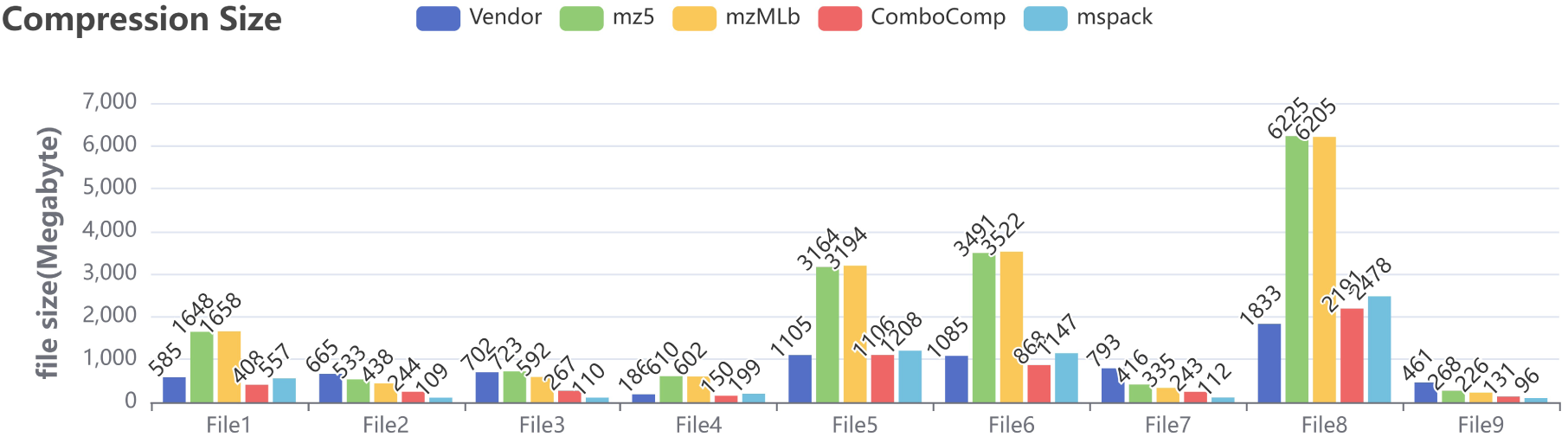
Size comparison with vendor format, mz5, mzML, mzMLb and mspack. mspack uses default loss parameters (–mz-fixed-abs, –int-log). For vendor format, mz5, mzML and mzMLb, ComboComp has great compression advantages. For mspack, the results show that ComboComp has better compression ratios in five of the nine files tested, while mspack achieves a more pronounced compression advantage in the remaining three files. It should be noted that mspack is a storage-oriented format, it does not support random file reading while Aird-ComboComp is a computationally oriented format that supports high-performance random file reading.

### ComboComp in MZmine3 and AirdPro

AirdPro supports manual selection of compression combinations for each dimension(see Figure 8). Users can choose the automatic decision mode to make automatic decisions. The decision parameters include compression ratio, compression time and decompression time. Users can freely adjust the target weight, and the combination generated by the decision maker will tend to the dimension with the largest weight.

Aird is also supported in MZMine3^21^(see Figure 9), which is a popular metabolomics data analysis software. Researchers can download mzmine3 or the source code from Github: https://github.com/mzmine/mzmine3.

**Figure 6:**
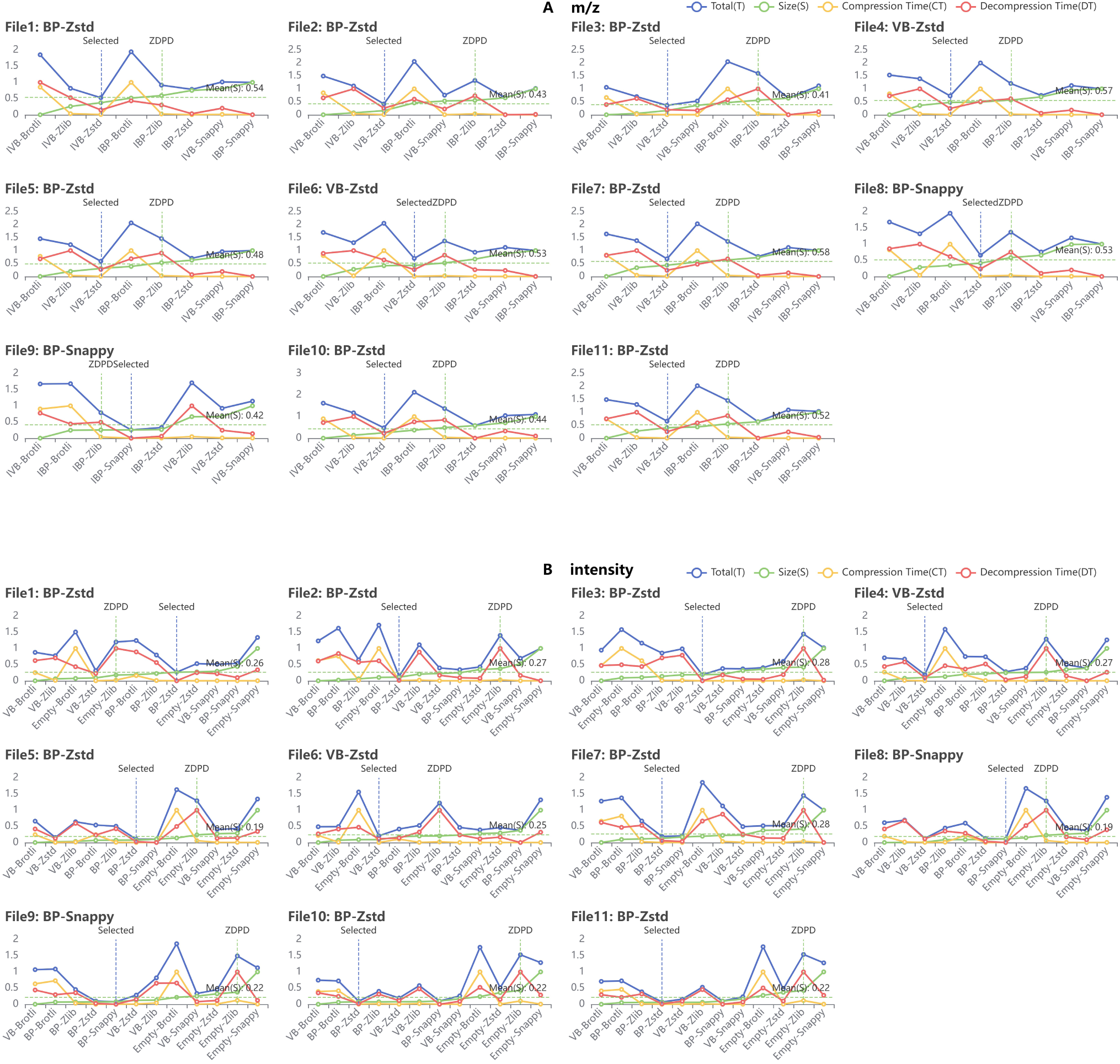
(A).Decision on *m/z*. The difference between the combination selected by the decider and the IVB+Zlib combination from ZDPD. (B).Decision on Intensity. The difference between the combination selected by the decider and the Empty+Zlib combination from ZDPD

**Figure 7:**
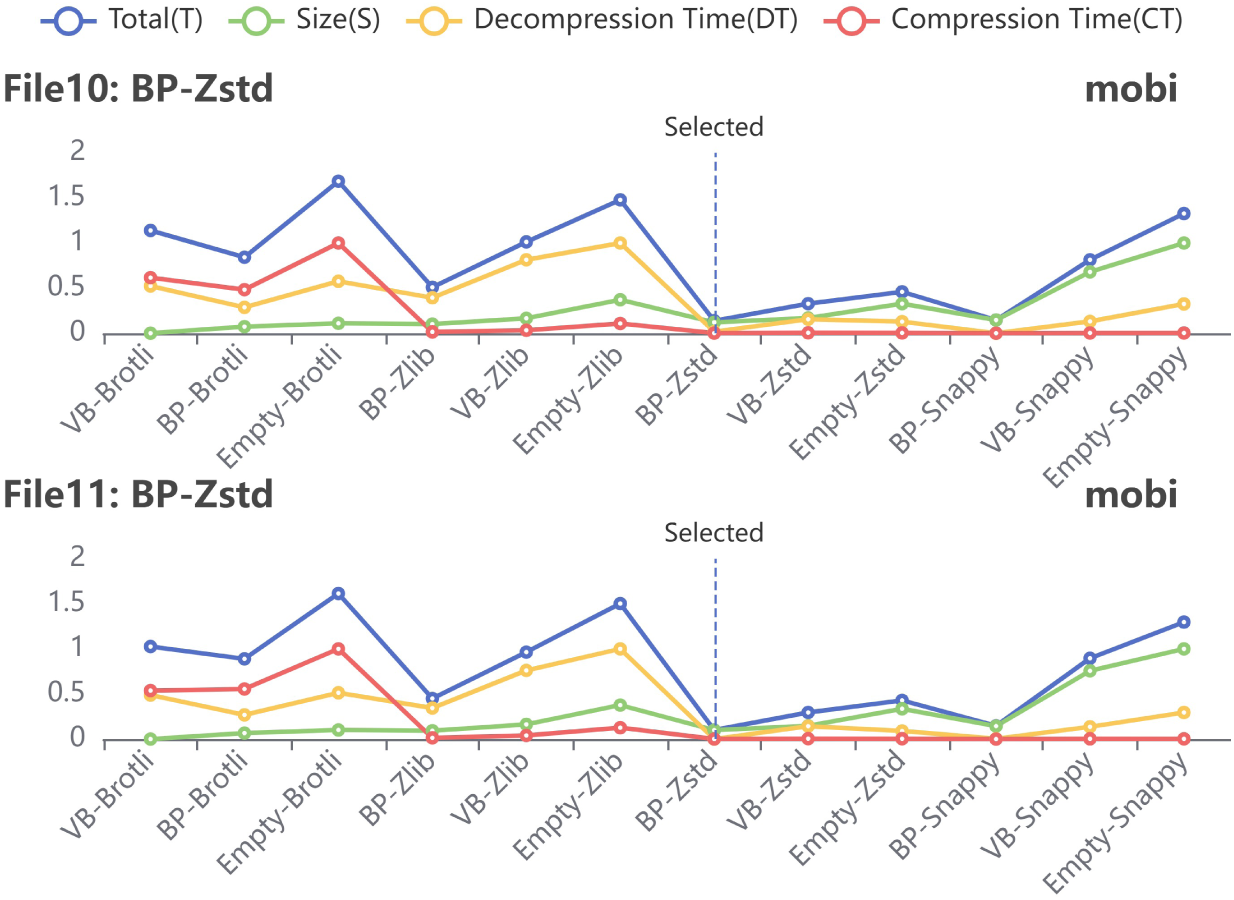
Decision on mobility. File10 and File11 have the mobility dimension in data. ComboComp shows excellent decision-making ability for ion mobility dimension as *m/z* and intensity. Among them, BP-Zstd delivers the best compression performance

**Figure 8:**
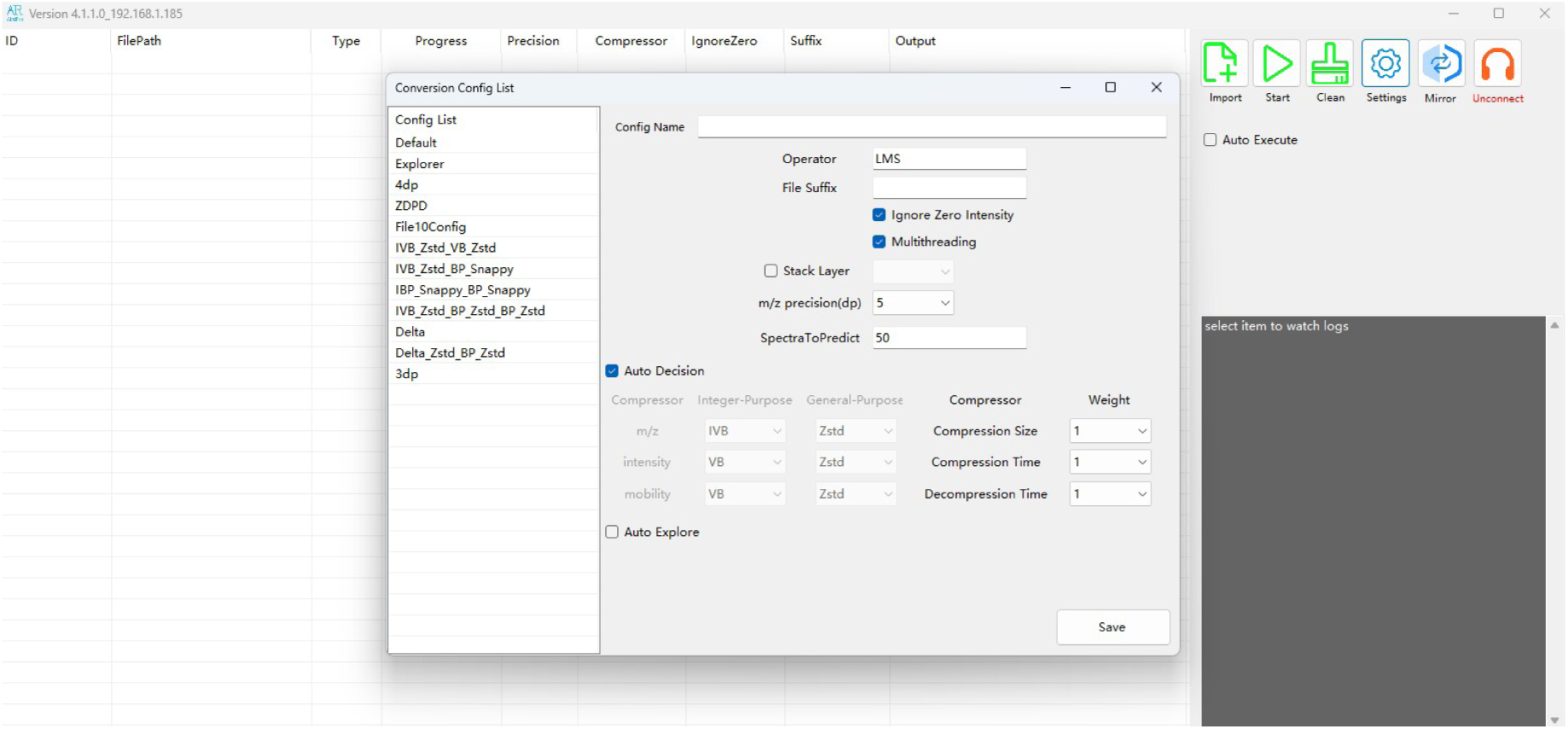
The main view of the AirdPro that supports for ComboComp

**Figure 9:**
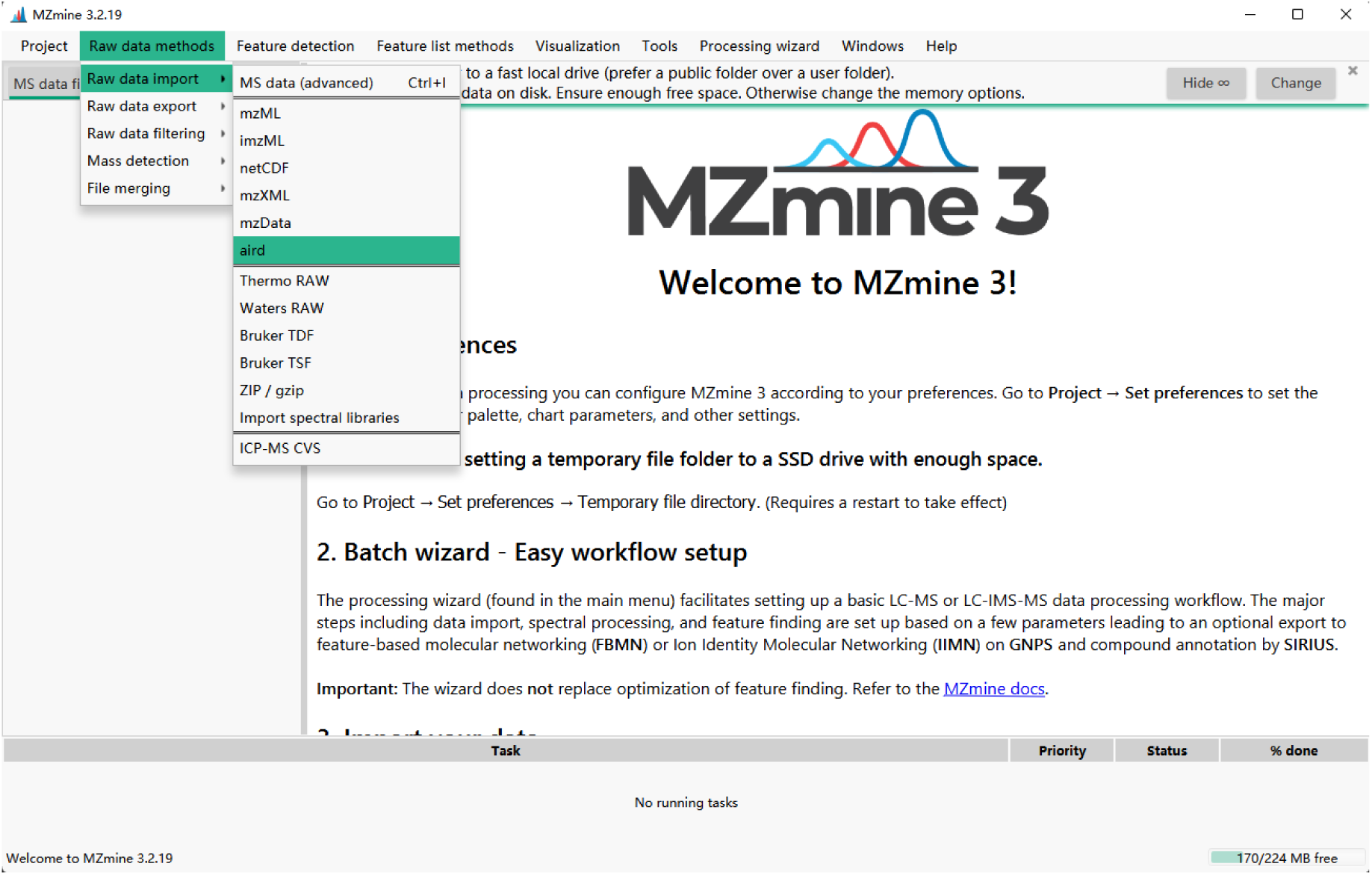
Aird support in MZmine3. Users can import aird files from the menu.

ComboComp is a new compression kernel for the Aird format. It implements high-performance compressions with random file access support. ComboComp provides a stable compression framework that compresses data in the three dimensions of *m/z*, intensity and ion mobility. It balances the decompression speed performance while considering the compression ratio, which greatly improves the format for downstream computing. The high compression rate can reduce the hardware requirement of mass spectrum data analysis, which can also effectively solve the high cost of cloud computing problem caused by the large volume and long queue of high-throughput MS data. MS data can even be analyzed in real time on IoT devices. We believe that ComboComp will find better compression kernels or compression combinations in the future to provide more guarantee for cloud computing or edge computing of MS data.

## Conclusions

Storing by column can significantly minimize the amount of data read from disk while computing XIC with a single *m/z*, and enhance search performance significantly. It also has the lowest CPU and memory consumption. Column storage loses its speed advantage when faced with hundreds or thousands of *m/z* as the data read steadily gets closer to reading the complete data file. Nevertheless, there are many situations in which it is sufficient to look through existing MS files just for several relevant chemical targets rather than fully analyze the entire file. Typically, this takes place during the data reanalysis stage. Aird-Slice provides a feasible solution for fast data search capability in MS data centers. When a new molecular target is hypothesized, the Aird-Slice can be used to quickly verify it in existing MS files. In addition to real-time search. Due to the fact that each column of data carries the whole signal of an *m/z*, Therefore, XIC calculation can be performed directly during data transmission. This also provides technical feasibility for streaming computing.

Mass spectrometer, as a technology that converts biological samples into digital samples, can effectively preserve valuable biological information. However, current algorithms and software are difficult to use all the signals in the MS file effectively. This means that current MS files may have new discoveries in the future. How to quickly search for the signal of the target molecule in an existing MS file has important biological significance. In addition, MS data analysis using Aird-Slice has minimal hardware requirements and can be quickly applied in IoT devices. This makes decentralized MS data analysis possible.

The following abbreviations are used in this manuscript:

## Abbreviations

PSI: Proteomics Standards Initiative
CV: Controlled Vocabular
MS: Mass spectrometry
XML: Extensible markup language
SDK: Software Development Kit
ZDPD: Zlib-Diff-PforDelta
TIMS: Trapped ion mobility spectrometry
m/z: Mass to charge ratio
HDF5: Hierarchical Data Format version 5
JSON: JavaScript Object Notation
SIMD: Single instruction multiple data
DDA: Data dependent acquisition
DIA: Data independent acquisition
IMS: Ion mobility spectrometry
XIC/EIC: Extracted ion chromatogram
PASEF: Parallel accumulation serial fragmentation
dp: Decimal places
GB: Gigabyte
MB: Megabyte
BP: Binary Packing
VB: Variable Byte
IVB: Variable Byte with delta encoding
IBP: Binary Packing with delta encoding
SSE: streaming SIMD extension

## Acknowledgement

We thank Jiawei Wang and Xixi Wang for testing ComboComp.

